# *Scn2a* deletion disrupts oligodendroglia function: Implication for myelination, neural circuitry, and auditory hypersensitivity in ASD

**DOI:** 10.1101/2024.04.15.589242

**Authors:** Han-Gyu Bae, Wan-Chen Wu, Kaila Nip, Elizabeth Gould, Jun Hee Kim

**Affiliations:** Kresge Hearing Research Institute, Department of Otolaryngology Head and Neck Surgery, University of Michigan, Ann Arbor, MI; Department of Cell and Developmental Biology, University of Michigan, Ann Arbor, MI; Department of Cellular and Integrative Physiology, University of Texas Health Science Center, San Antonio, TX

## Abstract

Autism spectrum disorder (ASD) is characterized by a complex etiology, with genetic determinants significantly influencing its manifestation. Among these, the *Scn2a* gene emerges as a pivotal player, crucially involved in both glial and neuronal functionality. This study elucidates the underexplored roles of *Scn2a* in oligodendrocytes, and its subsequent impact on myelination and auditory neural processes. The results reveal a nuanced interplay between oligodendrocytes and axons, where *Scn2a* deletion causes alterations in the intricate process of myelination. This disruption, in turn, instigates changes in axonal properties and neuronal activities at the single cell level. Furthermore, oligodendrocyte-specific *Scn2a* deletion compromises the integrity of neural circuitry within auditory pathways, leading to auditory hypersensitivity—a common sensory abnormality observed in ASD. Through transcriptional profiling, we identified alterations in the expression of myelin-associated genes, highlighting the cellular consequences engendered by *Scn2a* deletion. In summary, the findings provide unprecedented insights into the pathway from *Scn2a* deletion in oligodendrocytes to sensory abnormalities in ASD, underscoring the integral role of *Scn2a*-mediated myelination in auditory responses. This research thereby provides novel insights into the intricate tapestry of genetic and cellular interactions inherent in ASD.

## Introduction

Autism spectrum disorder (ASD) is an early neurodevelopmental disorder with complex neurological conditions. A prevailing hypothesis suggests that ASD emanates from alterations in brain connectivity(*1–3*). Myelination is critical for brain connectivity and temporal processing in the developing brain by coordinating effective axonal conduction and neurotransmission(*4–6*). Recent studies using magnetic resonance imaging (MRI) and genetic analysis have shown atypical white matter development and myelination patterns in individuals with ASD and animal models (*7, 8*). However, the specific impact of myelin alternations on neural connectivity, communication, and ultimately behavioral manifestations is not fully delineated. Notably, white matter integrity and myelin thickness, altered in specific brain regions of humans with ASD, are associated with sensory processing(*9, 10*). Auditory processing abnormalities including auditory hypersensitivity, difficulty listening with background noise, and problems encoding speech sounds are well documented sensory processing abnormalities in individual with ASD(*9*). The auditory hypersensitivity, as an excessive or abnormal response to auditory stimuli, may arise from alterations in sensory gain, neuronal activity, and excitation/inhibition (E/I) balance in the auditory circuitry, potentially linked to abnormalities in myelination(*11–13*).

Oligodendrocytes (OLs), myelin producing glia, play a more complex role than just facilitators of myelination; immature OLs monitor neuronal activities and control their proliferation and differentiation(*14, 15*). Moreover, OLs engage in dynamic interaction with neurons, contributing to activity-dependent myelination, and consequently neural circuit plasticity(*16–18*). Thus, OL dysfunctions can impact the neuron-glia interaction and ion channel expression along the axon, influencing neural plasticity(*5, 19*). However, the impact of changes in OL physiological properties on myelination and neuronal activity at the single-cell level, and the subsequent effects on auditory processing within the context of ASD, remains unexplored.

The gene, *Scn2a*, encoding the alpha subunit of the voltage-gated Na^+^ channel 1.2 (Na_v_1.2), is highly linked to neurodevelopmental disorders including ASD(*20–22*). Loss-of-function mutations in *Scn2a* impair dendritic excitability, leading to synaptic dysfunction and behavioral deficits(*22–24*). While *Scn2a* expression in non-neuronal cells, notably oligodendroglia(*25–27*), has been evident, its role in OL development and function remains to be elucidated. Notably, a transcriptional profile of OL lineage cells from mouse brain showed *Scn2a* expression in OLs and the highest levels of its expression are in newly formed OLs, an immature OL population(*25, 27*). A subpopulation of immature OLs express Na_v_1.2 channels, which mediated oligodendroglia spikes in the auditory brainstem and cerebellum, indicating *Scn2a* is important for electrical excitability in immature OLs(*26–28*). Here, we investigated how *Scn2a* deletion specifically in OLs impacts the interplay between myelination, neuronal activity, and auditory hypersensitivity that could further elucidate the neurological underpinnings of sensory processing disorders in ASD.

## Results

### *Scn2a* deletion in immature OLs altered gene expression pattern in mature OLs in *Scn2a* cKO mice

To understand the impact of *Scn2a* deletion in immature OLs on OL development and myelination, we characterized the single cell transcriptional profiles of mouse auditory brainstem using single nucleus RNA sequencing (snRNA-seq), which used isolated nuclei from the brainstem of control mice (*Scn2a*^fl/fl^) and *Scn2a* cKO mice (*Pdgfra*^CreERT^; *Scn2a*^fl/fl^) at P21-22. After the exclusion of low-quality cells, we obtained a total of 3083 reliable nuclei for the analysis: 989 nuclei from the control and 2094 nuclei from *Scn2a* cKO mice. Shared Nearest Neighbor (SNN) clustering identified 16 distinct clusters on Uniform Manifold Approximation and Projection (UMAP) plot, including clusters of neurons (*Rbfox3* positive), astrocytes (*Aldh1l1* positive), microglia (*Itgam* positive), and OL lineage cells (*Pdgfra* and *Mog* positive). Specifically, OL lineage cells consisted of five clusters: Oligodendrocyte precursor cells (OPCs), Differentiating OL1 (Diff. OL1), Differentiating OL2 (Diff. OL2), Premature OL, and Mature OL (**Fig. 1B**, **Fig. S1**).

**Fig. 1.**
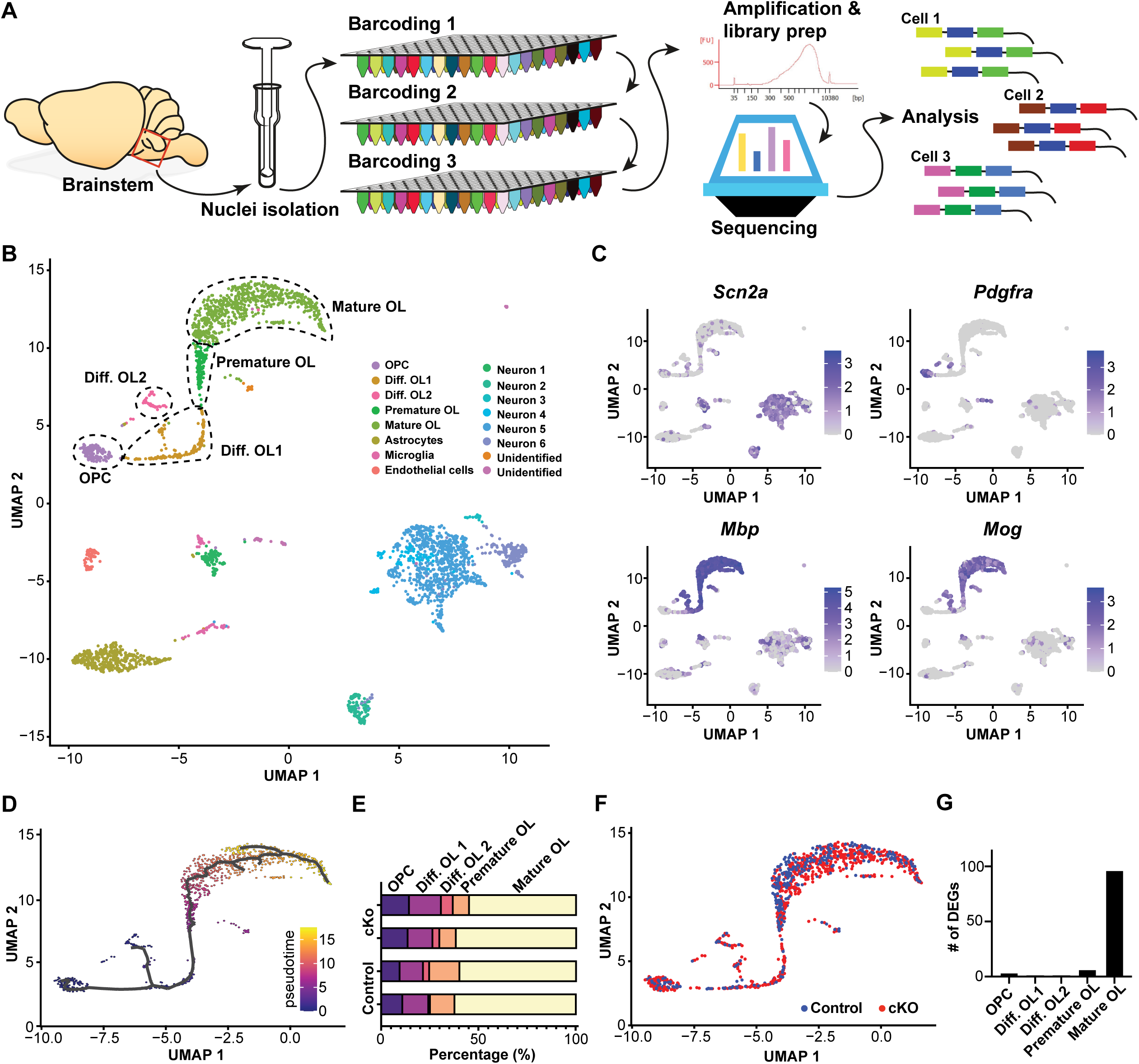
Single cell transcriptional profiles of mouse brainstems with OL-specific deletion of *Scn2a*. **A.** Illustration for single nuclei RNA sequencing (snRNA-seq) of the mouse brainstem. Control and *Scn2a* cKO mice brainstems (*n* = 4) were analyzed with snRNA-seq. **B.** UMAP plots of snRNA-seq dataset. Total 16 distinct clusters were detected. Upper left clusters were identified as OL lineage cells, indicated by broken-line circles. OPC, Differentiating OL1 (Diff. OL1), Differentiating OL2 (Diff. OL2), Premature OL, and Mature OL. Other clusters represent astrocyte, microglia, and neurons based on specific gene expression. **C.** Expression pattern of cell-type specific genes such as *Scn2a, Pdgfra, Mbp* and *Mog*, based on the scaled gene expression level (graded color indicators, right). **D.** The distribution of single cells colored according to the five OL lineage states identified by trajectory analysis, from OPCs to mature OLs. **E.** Percentage (%) of each cluster (from OPC to Mature OL) in total OLs in individual control and cKO mice. **F.** UMAP plots of snRNA-seq for OL lineage cells from control (blue dots) and cKO (red dots). **G.** The number of DEGs for each OL lineage. There is a distinct difference in gene expression in mature OLs between control and cKO mice.

*Scn2a* was strongly expressed in neurons. Notably, it was also detected across the OL lineage, supporting *Scn2a* expression and function in the OL lineage. To identify the OL lineage, we analyzed OL-specific genes, such as *Pdgfra*, *Mbp* and *Mog*, expressed in OPCs and mature OLs, respectively (**Fig. 1C**). Given the dynamic nature of OL differentiation, we utilized trajectory analysis to examine this continuous differentiation. The result of trajectory analysis from those populations displayed a narrow differentiation path connecting from OPCs to mature OLs (**Fig. 1D**), that similar to patterns in a previously reported study(*25*). The result indicates that OL development follows a clearly defined trajectory, which was outlined by genetic markers specific to the OL lineage (e.g., *Pdgfra* for OPCs; *Mbp* and *Mog* for mature OLs). When comparing the populations of each cluster between the control and *Scn2a* cKO mice, there was no statistically significant difference in the proportion of OL lineage cells. However, a notable trend towards a reduction in the mature OL population was observed in the *Scn2a* cKO mice, whereas OPCs and differentiating OLs exhibited a slight increase (**Fig. 1E**). This result supports a previous study demonstrating that OL-*Scn2a* contributes to OL differentiation(*27*). Additionally, when OL clusters from each genotype were visualized with different colors in the UMAP scatter plot, a distinct difference was observed in mature OLs (**Fig. 1F**). To quantify these differences further, we identified genes exhibiting significant differential expression between control and *Scn2a* cKO mice (adjusted *p* with Bonferroni correction < 0.05). Notably, 96 differentially expressed genes (DEGs) were detected in mature OLs between control and *Scn2a* cKO mice, while only 3, 1, 1, and 6 DEGs in OPC, differentiating OL, differentiating OL2, and premature OL groups, respectively, were identified (**Fig. 1G**, **Table S1-5**). This result reveals significant alterations in gene expression patterns within mature OLs, suggesting the cascading implications of *Scn2a* deletion in the orchestration of gene expression within mature OLs.

### Myelin related genes were significantly downregulated in mature OL from *Scn2a* cKO mice

Furthermore, to examine the expression level of myelin-associated genes in mature OLs, we employed AmiGo2, a gene ontology data base, to pick genes for comparison. Using the keyword "myelin," we identified a total of 391 genes and their expression levels were compared in control and *Scn2a* cKO mice. Among these myelin-related genes, 33 DEGs were identified (*p* < 0.05). For example, *Mobp*, a gene encoding myelin associated oligodendrocyte basic protein; MOBP, was significantly decreased in *Scn2a* cKO mice (**Fig. 2A**). The String analysis revealed that 30 genes out of 33 DEGs have a significant interconnection and can be functionally characterized into four main categories: cell skeletal structure, cell junction, membrane, and ion channel (**Fig. 2B**). In addition, the most significant down regulated genes; *Mobp*(*29*), *Ncam1*(*30*), *Ptn*(*31*), *Gnao1*(*32*), *Mtmr2*(*33*), *Abca2*(*34*), *Arhgef10*(*35*) and *Mpdz*(*36*) are known to be related to myelination, cell growth, and neurological diseases (**Fig. 2C**). Taken together, the transcriptional profiles highlight a pronounced differential expression of myelin-associated genes in mature OLs from *Scn2a* cKO mice.

**Fig. 2.**
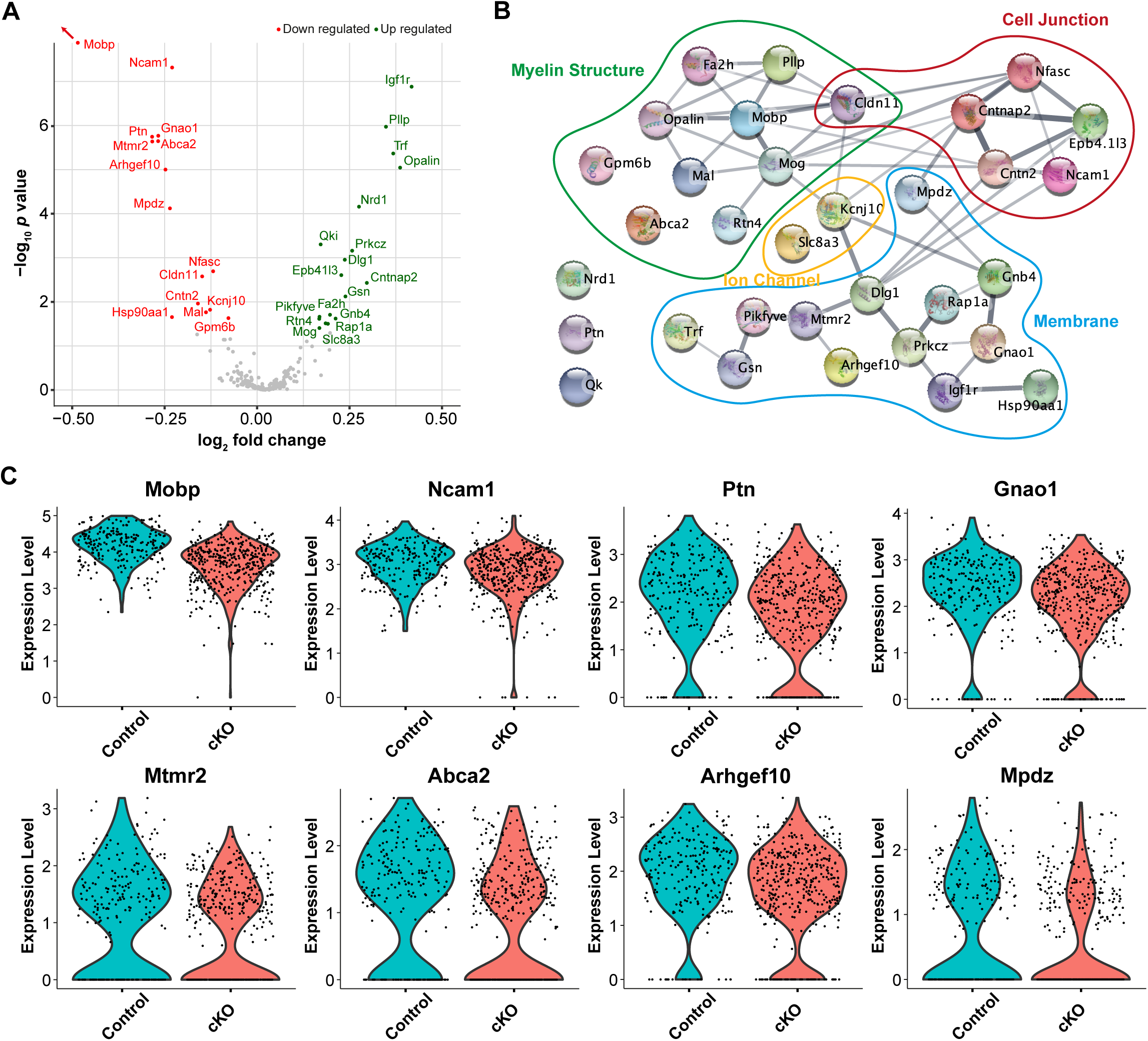
Genes related to myelination are significantly decreased in mature OLs from *Scn2a* cKO mice. **A.** Volcano plot of differentially expressed myelin related genes (391 genes) in control and cKO mice. **B.** 33 genes showed significant differential expression (*p* < 0.05) and interconnection between those genes was detected in Sting analysis. Those genes were related to cellular structure, cell junction, membrane protein signaling (e.g. G-protein mediated signaling). **C.** Violin plots of top eight genes being downregulated in *Scn2a* cKO mice (red); *Mobp, Ncam1, Ptn, Gnao1, Mtmr2, Abca2, Arhgef10* and *Mpdz*, of which deficiency is known to contribute to abnormal myelination.

### Loss of *Scn2a* impairs myelination and alters axon caliber size in the auditory brainstem during postnatal development

Following the gene expression analysis, we assessed the structural aspects of myelinated axons using transmission electron microscopy (TEM, **Fig. 3A**). This evaluation focused on determining myelin thickness and axon caliber size within axon bundles located in the medical nucleus of trapezoid body (MNTB) in the auditory brainstem. Ultrastructural analysis revealed significant myelin deficits in *Scn2a* cKO mice, as evidenced by an elevated *g*-ratio providing critical insights into the intricate alterations occurring at the structural level (**Fig. 3B**). The *g*-ratio is an indicator of myelin thickness measured by the inner radius (r) divided by the outer radius (R) of axon in the corresponding axon bundles. To examine alteration in distribution patter of the *g*-ratio, linear regression analysis was used in which slope and y-intercept were compared between groups. While the regression slopes of control and *Scn2a* cKO were comparable (0.0385 in control mice vs 0.0399 in *Scn2a* cKO mice, *p* = 0.6295, linear regression comparison), y-intercept was significantly altered (0.7616 in control mice vs 0.7766 in *Scn2a* cKO mice, *p* < 0.0001, linear regression comparison). Axon calibers showed a larger inner diameter in *Scn2a* cKO compared to control (1.39 ± 0.03 µm, n = 1154 axons in 10 control mice vs 1.48 ± 0.03 µm, n = 786 axons in 4 *Scn2a* cKO mice, *p* < 0.0001, Mann-Whitney *U*-test, **Fig. 3C**). The right shift of the cumulative frequency curve indicates an increase in axon caliber size in *Scn2a* cKO compared to control (**Fig. 3C**). Intriguingly, the *g*-ratio was significantly increased in *Scn2a* cKO, indicating a decrease in myelin thickness of axons in the MNTB area (0.82 ± 0.002, n = 1154 axons in 10 control vs 0.84 ± 0.002, n = 786 axons in 4 *Scn2a* cKO mice, *p* < 0.0001, Mann-Whitney *U*-test, **Fig. 3D**). In addition, there was also a right shift in the cumulative frequency distribution of *g*-ratio in *Scn2a* cKO, indicating *g*-ratio was larger in *Scn2a* cKO across examined axons. The results suggest that the targeted deletion of *Scn2a* in immature OLs detrimentally affects myelination, concurrently altering the axon caliber size within the auditory brainstem. Notably, the myelin deficits identified in *Scn2a* cKO mice were consistent with significant alterations in the expression of myelin-related genes in mature OLs from snRNA-seq data (**Fig. 2**).

**Fig. 3.**
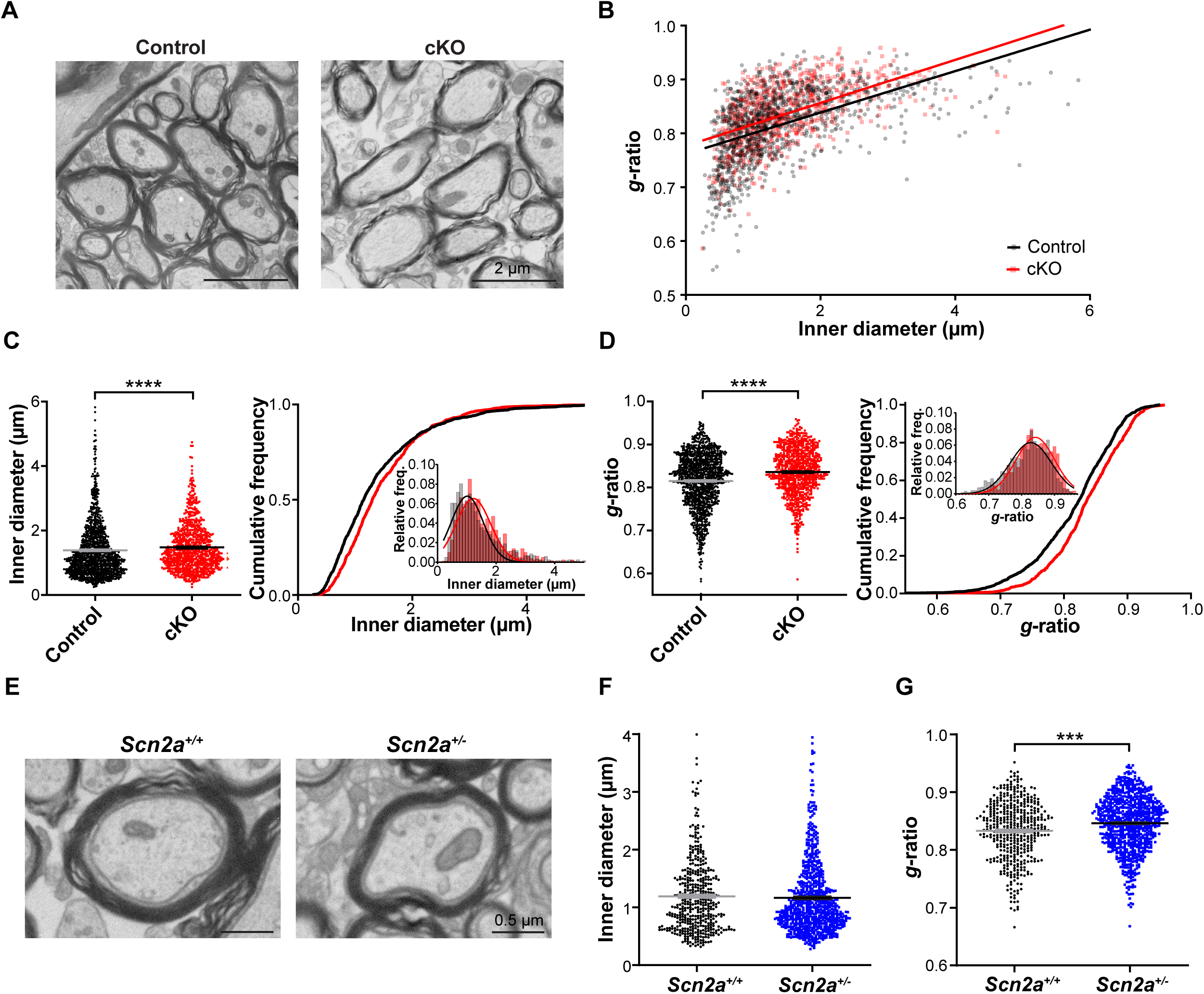
Loss of *Scn2a* in OLs increases axon caliber and decreases myelin thickness in mouse auditory brainstem. **A.** Representative TEM images of axons in the MNTB from control and *Scn2a* cKO mice at P25. **B.** Scatter plot of *g*-ratio values plotted as a function of corresponding axon inner diameter of individual axons. Datapoints from individual axons and best fit lines are shown in black for control and red for *Scn2a* cKO. **C.** Summary of inner diameter of individual axons (left) and the respective cumulative frequency (right). Inset, the histogram distribution of inner diameter. **D.** Summary of *g*-ratio of individual axons and the respective cumulative frequency. Inset, the histogram of distribution of *g*-ratio. **E.** Representative TEM images of an axon in the MNTB from conventional loss-of-function *Scn2a* heterozygous (*Scn2a*^+/-^) and wildtype (*Scn2a*^+/+^) mice. **F.** Summary of inner diameter of individual axons from *Scn2a*^+/+^ and *Scn2a*^+/-^ mice. **G.** *g*-ratio of individual axons from *Scn2a*^+/+^ and *Scn2a*^+/-^ mice. Values are shown as mean ± s.e.m., *** *p* < 0.001, **** *p* < 0.0001.

For further validation, we tested effect of *Scn2a* deletion on myelination in different mouse models. Using conventional *Scn2a* heterozygous mice with a loss-of-function mutation in *Scn2a* (*Scn2a*^+/−^)(*22, 37*), we examined the effect of *Scn2a* reduction on myelination. We quantified the myelin sheath thickness and axon caliber size of axons in the MNTB. The *g*-ratio was significantly higher in *Scn2a*^+/–^ mice compared to wildtype mice (*Scn2a*^+/+^) without a significant difference in axon diameter (*g*-ratio, 0.83 ± 0.003, n = 439 axons in 5 *Scn2a*^+/+^ mice vs 0.85 ± 0.002, n = 768 axons in 4 *Scn2a*^+/–^ mice, *p* < 0.0001, Mann-Whitney *U*-test, **Fig. 3E-G**). The result suggests that reduced *Scn2a* affects myelin development in the auditory brainstem during postnatal development. In addition, another OL-specific *Scn2a* cKO mice, in which *Scn2a* was removed from Sox10 expressing OLs, targeting all OL lineage cells (*Sox10*^CreER^; *Scn2a*^fl/fl^)(*38*), also displayed similar alterations in myelin and axon caliber size (**Fig. S2**). The observed myelin deficits in *Scn2a* cKO mice can be attributed to both impaired OL development and alterations in myelin-associated genes. A reduction in the differentiation of OPCs to OLs, evidenced by a decreased count of CC1+ OLs in *Scn2a* cKO mice, further supports these findings(*27*). Together, these results collectively underscore the pivotal role of *Scn2a* in myelination and the maintenance of axonal integrity in the auditory brainstem.

### OL-specific loss of *Scn2a* alters myelinated segments at distal axons in the MNTB

Given the significant alterations in myelination, we postulated that myelin changes might manifest in broader structural and functional aberrations within the neural circuitry. During development, immature OLs play a pivotal role by interacting with axons, which in turn determines the length of myelinated axon segments and nodes (*5*). To elucidate the consequences of *Scn2a* deletion in OLs on these interactions, we examined the long-myelinated axon extending from the cochlear nucleus to the MNTB and its terminal, the calyx of Held. We quantified the length of myelinated axon segments. The distal axons in the MNTB are fully myelinated and each myelinated segment can be visualized by myelin proteolipid protein (PLP) flanked by two sodium channel clusters (Pan Na_v_) denoting nodes of Ranvier (**Fig. 4A**). Thus, we assessed the length of myelinated segments from the internode length by measuring the distance between two Na_v_ clusters (**Fig. 4B**). A structural analysis of the calyx axons showed a significant reduction in internodal length adjacent to the heminode at the distal axon in *Scn2a* cKO mice (42.01 ± 2.31 μm, n = 24 axons from 5 control mice vs 32.46 ± 1.83 μm, n = 46 axons from 5 cKO mice, *p* = 0.0024, unpaired *t*-test, **Fig. 4C**). However, no significant difference was observed in the internodal length from node to node (52.78 ± 4.74 μm, n = 12 axons from 5 control mice vs 55.56 ± 4.33 μm, n = 16 axons from 5 cKO mice, *p* = 0.6703, unpaired *t*-test, **Fig. 4C**). The reduced length of the last myelinated segment length in *Scn2a* cKO suggests that myelinated segments near the nerve terminal might be underdeveloped in *Scn2a* cKO mice(*39*).

**Fig. 4.**
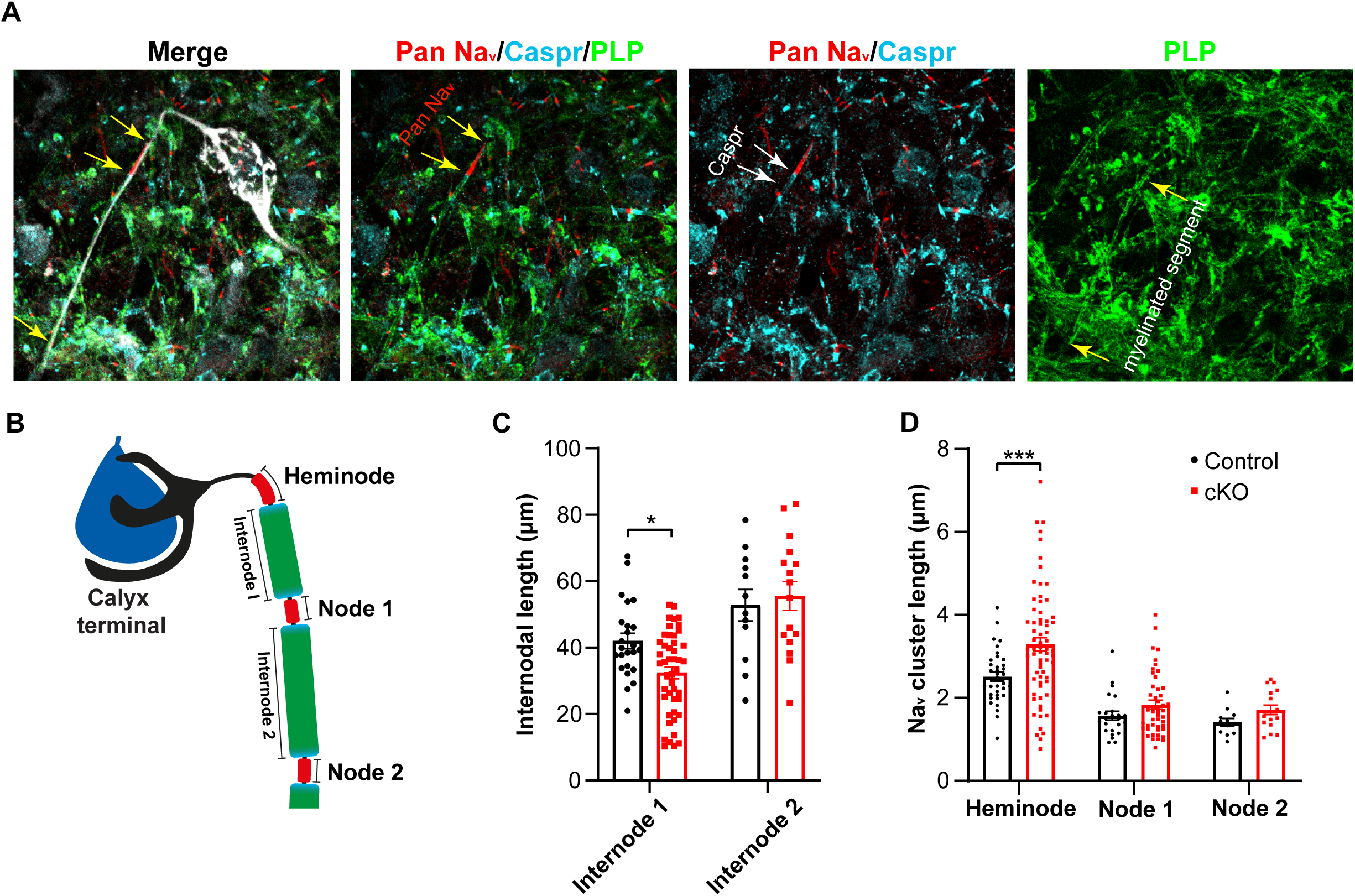
OL dysfunctions impact axonal integrity along the distal axon in the developing brainstem. **A.** The calyx of Held terminal was dye-filled during whole-cell recording (Lucifer yellow) and subsequently the MNTB was immunostained with antibodies against myelin proteolipid protein (PLP, green) Pan Nav (red, yellow arrows), and Caspr (cyan, white arrows). **B.** Illustration of calyx of Held axon showing the heminode, nodes, and internodes (myelinated segments). **C.** Summary of internodal length, which was measured by distance from the heminode to node 1 (internode 1) and node 1 to node 2 (internode 2) along the distal axon. **D.** The length of Nav clusters (Pan Na_v_) at heminode, node1, and node 2. Values are shown as mean ± s.e.m., * *p* < 0.05, *** *p* < 0.001.

Furthermore, to determine if the thinner and shorter myelin alters ion channel distribution along the distal axon in *Scn2a* cKO mice, we examined the patterns of voltage-gated sodium channel (Na_v_) expression along the calyx of Held axon. Intriguingly, we found a dispersed Na_v_ channel expression at heminodes in *Scn2a* cKO mice (2.51 ± 0.11 μm, n = 35 axons from 5 control mice vs 3.29 ± 0.17 μm, n = 67 axons from 7 cKO mice, *p* = 0.0018, unpaired *t*-test, **Fig. 4D**). Dispersed Na_v_ channels at the heminode have also been previously observed in dysmyelinated axon terminals(*5*). Nevertheless, Na_v_ clusters at nodes did not show any significant alteration (unpaired *t*-test, *p* = 0.1209 for node 1 and *p* = 0.616 for node 2, **Fig. 4D**). This result highlights the pivotal role of *Scn2a* in OL development, emphasizing its influence on axon-OL interactions and Na_v_ channel distribution at the distal axons in synapse-rich areas of the auditory brainstem during postnatal development.

### Conduction failures impaired the reliability and fidelity of high frequency spikes at the nerve terminal in *Scn2a* cKO mice

Alterations in Na_v_ channel clustering at the heminode have been shown to impair the fidelity of presynaptic action potentials (APs), leading to an increased failure rate during high-frequency spiking in the developing brainstem.(*5, 40*) To understand the implications of thinner myelin sheath, larger axon, and ion channel expression alterations at the distal axon observed in *Scn2a* cKO, we applied electrical stimuli along the calyx axons and evaluated the fidelity of APs at the nerve terminal (**Fig. 5A**). In control, the calyx terminal efficiently followed high-frequency stimulation and demonstrated spikes without failures at both 50 Hz and 100 Hz. Conversely, the calyx terminal in *Scn2a* cKO mice exhibited AP failures at 100 Hz and had drastically more failures at higher frequencies (**Fig. 5B, C**). For example, in response to 200 Hz stimulation, 28% of axons in the *Scn2a* cKO displayed failures compared to only 18% in control mice. Furthermore, violin plots revealed a larger proportion of axons with over 80% failure at higher frequencies (300-500 Hz) in *Scn2a* cKO mice while no axons in control mice showed failure rate above 80% in the same frequency range (**Fig. 5C**). Thus, myelin defects and disruption of Na_v_ channel clustering caused by *Scn2a* loss detrimentally affect the reliability and temporal fidelity of spikes at the nerve terminal.

**Fig. 5.**
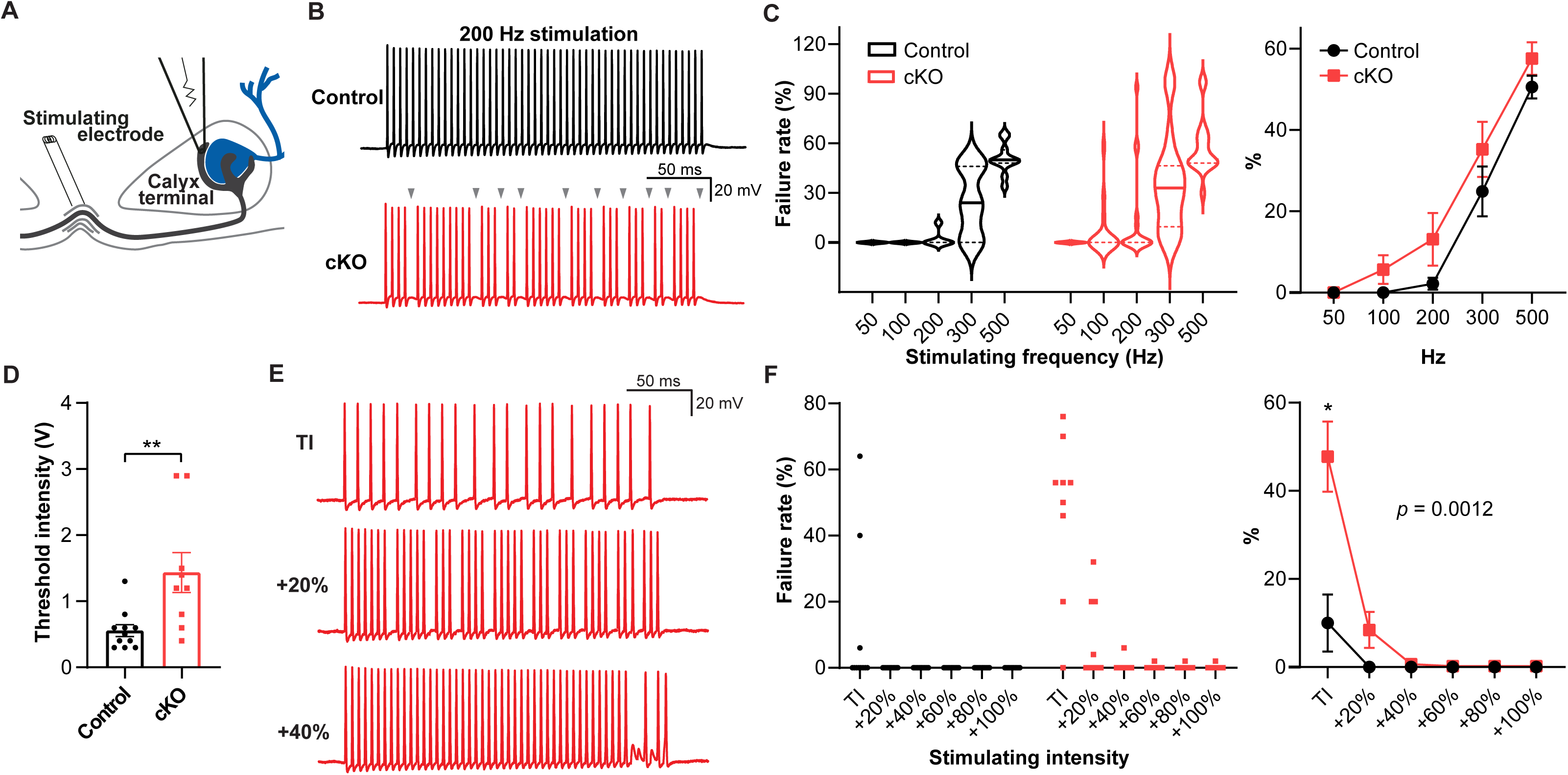
The conduction and fidelity of spikes at the nerve terminal are impaired in *Scn2a* cKO mice. **A.** Illustration of the calyx of Held recording with a bipolar stimulator placed in the midline of the brainstem. **B.** Representative traces of presynaptic AP train stimulated at 200 Hz from control (black) and *Scn2a* cKO (red) mice. Grey arrowheads indicate AP failure during the train in *Scn2a* cKO mice. **C.** Left, violin plots of AP failure rate from control and *Scn2a* cKO mice stimulated at varied frequencies (from 50 Hz to 500 Hz). Solid and dashed lines represent median and first and third quartile, respectively. Right, average AP failure rate at varied stimulating frequencies. **D.** The average stimulating intensity to initiate a single AP (Threshold intensity, TI) from control and *Scn2a* cKO. **E.** 200 Hz AP train from *Scn2a* cKO stimulated with TI, 20% increment from TI, and 40% increment from TI. Note that increasing stimulation intensity reduced AP failures. **F.** Left, 200 Hz AP failure rate reduced by incremental stimulating intensities in control and *Scn2a* cKO. Circles and squares indicate individual datapoint (n = 11 cells in control vs n = 18 cells in cKO). Right, summary from the left showing average failure rate from incremental stimulating intensities (mean ± s.e.m.). * *p* < 0.05, ** *p* < 0.01.

To discern whether AP failures at the nerve terminal are caused by conduction failures throughout the distal axon or failure to evoke AP, we tested if increasing the stimulus intensity could recover AP failures. First, we determined the minimum stimulating intensity (threshold intensity, TI) necessary to trigger a single AP for each axon. We found that threshold intensity was significantly higher in *Scn2a* cKO than in control (0.55 ± 0.09 V, n = 11 from 3 control mice vs 1.43 ± 0.30 V, n = 9 from 4 cKO mice, *p* = 0.0048, Mann-Whitney *U*-test, **Fig. 5D**). Next, we quantified failure rate in AP train (200 Hz stimulation) with incremental intensities (**Fig. 5E**). The calyx terminal from *Scn2a* cKO showed significantly higher failure rates at the TI (10 ± 6.49 %, n = 11 from 3 control mice vs 47.78 ± 7.95 %, n = 9 from 4 cKO mice, *p* = 0.0117, Two-way repeated measures ANOVA with Sidak correction, **Fig. 5F**) which was recovered by increased stimulating intensity. Interestingly, all axons in control with failures at TI were able to recover by TI + 20% intensity, whereas such intensity could only recover ∼56% of axons in *Scn2a* cKO. Most of the axons with failures in *Scn2a* cKO required at least TI + 40% to recover (*p* = 0.0012, Two-way repeated measures ANOVA, **Fig. 5F**). These findings were similarly observed in another OL-specific cKO mice (*Sox10*^CreER^; *Scn2a*^fl/fl^, **Fig. S3**). Taken together with EM and immunostaining analysis, these results suggest that a shortened distal internode and thinner myelin critically impair axon conduction throughout the distal axon and cause AP failures at the nerve terminal.

### The intrinsic excitability at the nerve terminal was impaired

Changes in myelination and Na_v_ channel clustering could lead to conduction failure as observed in *Scn2a* cKO. Notably, *Scn2a* cKO mice showed significantly higher stimulus intensity to generate AP compared to the control. This suggests that a stronger depolarization is required to evoke AP which implies changes in excitability. We therefore examined intrinsic excitability at the calyx terminal by recording presynaptic APs, evoked by depolarizing current injection. AP waveform analysis revealed that AP threshold was significantly higher in *Scn2a* cKO mice (-49.04 ± 0.76 mV, n = 32 from 17 control mice vs -46.88 ± 0.56 mV, n = 43 from 12 cKO mice, *p* = 0.0217, unpaired *t*-test, **Fig. 6A, B**), with no substantial changes in AP amplitude (77.58 ± 1.91 mV, n = 32 from 17 control mice vs 72.83 ± 1.95 mV, n = 43 from 12 cKO mice, *p* = 0.0912, unpaired *t*-test, **Fig. 6B**). The maximal dV/dt was lower in *Scn2a* cKO mice (468.10 ± 28.47 mV/ms, n = 32 from 17 control mice vs 411.80 ± 30.17 mV/ms, n = 43 from 12 cKO mice, *p* = 0.0282, Mann-Whitney *U*-test), indicating a slower AP rise time in *Scn2a* cKO mice than control. Furthermore, the minimal dV/dt was significantly reduced in *Scn2a* cKO mice (-359.80 ± 22.60 mV/ms, n = 28 from 17 control mice vs -292.9 ± 21.75 mV/ms, n = 36 from 12 ckO mice, *p* = 0.0061, Mann-Whitney *U*-test), indicating a slower repolarization. Other parameters in *Scn2a* cKO, despite not reaching statistical significance, showed trends of alteration: an increased rheobase current (84.79 ± 7.18 pA for control vs 103.60 ± 7.69 pA for cKO, *p* = 0.058, Mann-Whitney *U*-test) and a broadened AP width (0.28 ± 0.01 ms for control vs 0.30 ± 0.009 ms for cKO, *p* = 0.1437, unpaired *t*-test, **Fig. 6B**). An elevated threshold, a slower rising of the spike, and a slower repolarizing at the calyx terminal indicate that the nerve terminal has reduced excitability. To further support these findings, we also tested the intrinsic excitability by counting the number of APs evoked by incremental current injections. The input-output curves showed a distinct shift, demonstrating fewer APs per current step in *Scn2a* cKO (*p* = 0.0176, Two-way repeated measures ANOVA, **Fig. 6C**). Thus, the result suggests that alterations in Na_v_ channel expression around myelinated segments have a profound impact on presynaptic excitability in *Scn2a* cKO mice.

**Fig. 6.**
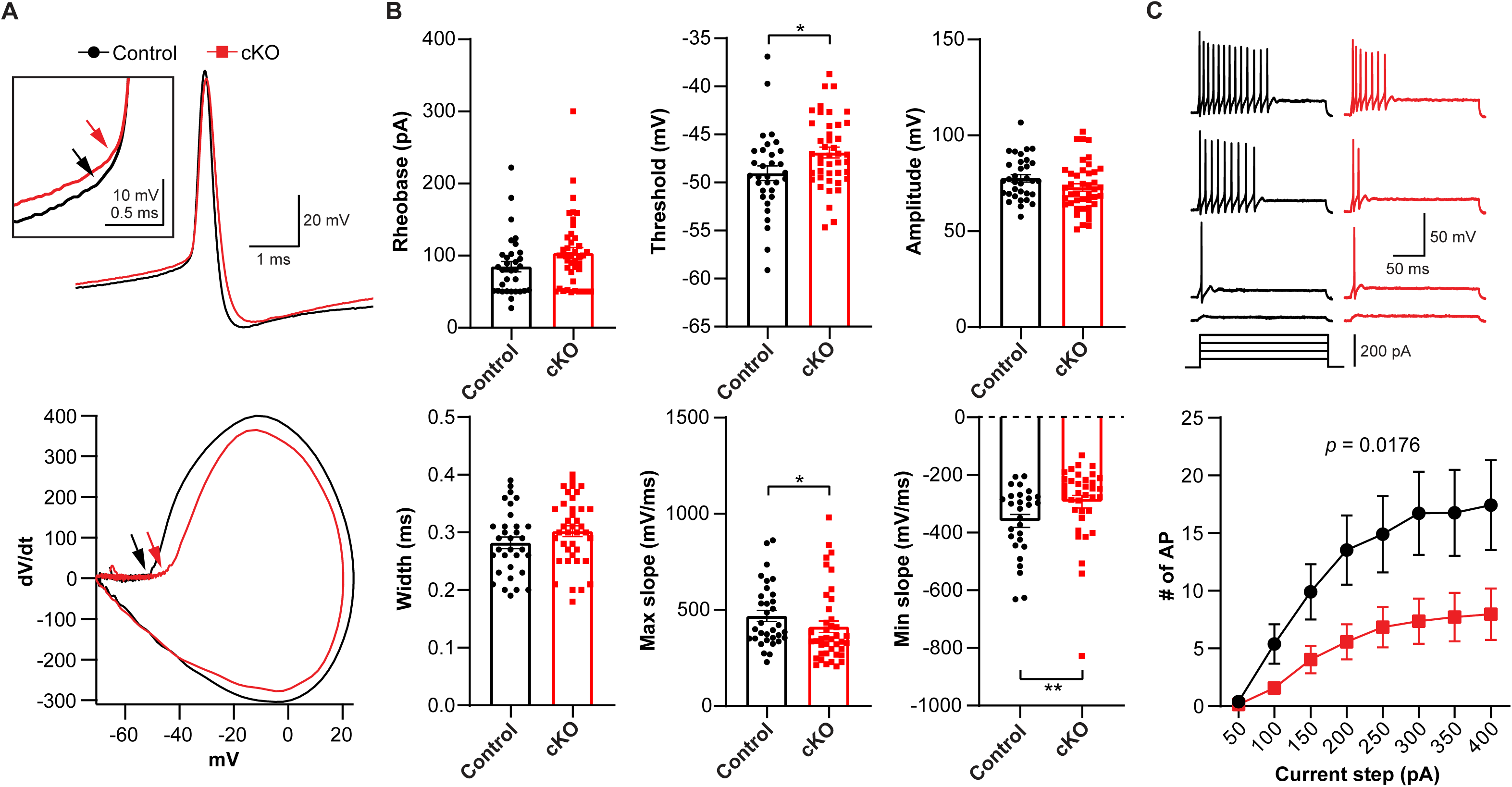
Intrinsic properties and excitability at the calyx terminal are altered in *Scn2a* cKO mice. **A.** Presynaptic APs elicited with a supra-threshold current injection and their corresponding phase plots (dV/dt against membrane potential) in control (black) and *Scn2a* cKO (red). Arrows indicate the threshold of AP. AP threshold was determined by the point where dV/dt exceeds 10 mV/ms and AP amplitude was calculated from the threshold to the AP peak in the phase plot. **B.** Summary of AP waveform analysis; the rheobase, threshold, amplitude, width, max dV/dt, and min dV/dt. Individual datapoints are displayed in each bar graph. **C.** Top, presynaptic APs elicited by step-like depolarizing current injection. Bottom, the number of AP in response to each current injection (200 ms, from 50 pA to 400 pA, mean ± s.e.m.). * *p* < 0.05, ** *p* < 0.01.

Intriguingly, another observation is that in ∼20% of presynaptic recordings from *Scn2a* cKO (14 of 45 cells), the calyx terminal displayed aberrant spikes or/and spontaneous spikes when the resting membrane potential was around -65 mV (**Fig. S4**). However, spontaneous firing and aberrant spikes have not been observed before in *ex vivo* recording from WT with the same experimental setting. The results demonstrated that myelin deficits at the distal axon caused by OL-*Scn2a* loss impaired conduction and the temporal fidelity of spikes at the nerve terminal, while generating asynchronous and abnormal spikes along the distal axon.

### *Scn2a* cKO mice exhibit auditory hypersensitivity without changes in peripheral function

Based on notable alteration in myelination and neuronal activity within the auditory pathways, we examined if these structural and functional anomalies have tangible behavioral repercussions in the *Scn2a* cKO mice. *SCN2A* is identified as a high-risk gene associated with ASD, wherein abnormal sensory processing including auditory hypersensitivity is frequently observed (*9, 41*). To ascertain the extent of auditory processing abnormalities in *Scn2a* cKO mice, we utilized *in vivo* auditory brainstem responses (ABRs) to assess the sum of evoked potential responding sound stimuli along the auditory pathway, particularly in the brainstem (**Fig. 7A**). There was no significant difference in ABR threshold (38.5 ± 2.1 dB of SPL, n = 46 control vs 34.0 ± 2.6 dB of SPL, n = 29 cKO mice, *p* = 0.1842, unpaired *t*-test, **Fig. 7B**) and the peak latencies between control and *Scn2a* cKO mice (*p* = 0.134, data not shown). However, the amplitudes of ABRs from *Scn2a* cKO mice significantly increased compared to those from control. The quantitative analysis of ABR amplitude in response to 85 dB SPL (click) showed that the amplitude of peak II (2.44 ± 0.166 µV, n = 39 control vs 3.30 ± 0.231 µV, n = 26 cKO, *p* = 0.0002, multiple comparison from two-way ANOVA with Bonferroni correction) and peak III (2.09 ± 0.129 µV in control vs 2.70 ± 0.149 µV in cKO, *p* = 0.0191) were significantly larger in *Scn2a* cKO mice compared to those of control. There was a trend toward an increase, but no significant changes were observed in peak I (1.04 ± 0.082 µV in control vs 1.41 ± 0.231 µV in cKO, *p* = 0.352, multiple comparison from two-way ANOVA with Bonferroni correction), peak IV (0.72 ± 0.07 µV in control vs 1.11 ± 0.14 µV in cKO, *p* = 0.3248, multiple comparison from two-way ANOVA with Bonferroni correction) and peak V (1.00 ± 0.094 µV in control vs 1.26 ± 0.166 µV in cKO, multiple t test, *p* >0.9999, multiple comparison from two-way ANOVA with Bonferroni correction, **Fig. 7C**). The increased amplitude of ABRs suggested that *Scn2a* cKO mice exhibit the hypersensitivity to sound. To evaluate whether there were alterations in central gain, we calculated the ratio between peak amplitudes II to IV relative to peak I. Notably, our analysis indicated a prevalent trend where the ratios of peak II to peak I and peak III to peak I were larger in *Scn2a* cKO mice, despite the absence of statistical significance (*p* > 0.05, two-way ANOVA, **Fig. 7D**). We examined the possibility if the hypersensitivity could have originated from alterations in peripheral hearing sensitivity. There was no significant change in ABR threshold, peak I amplitude, and Distortion Product Otoacoustic Emissions (DPOAE, **Fig. 7E**), demonstrating intact peripheral hearing. Therefore, the results suggest hypersensitivity observed in *Scn2a* cKO mice is predominantly induced by changes within central auditory brain circuits.

**Fig. 7.**
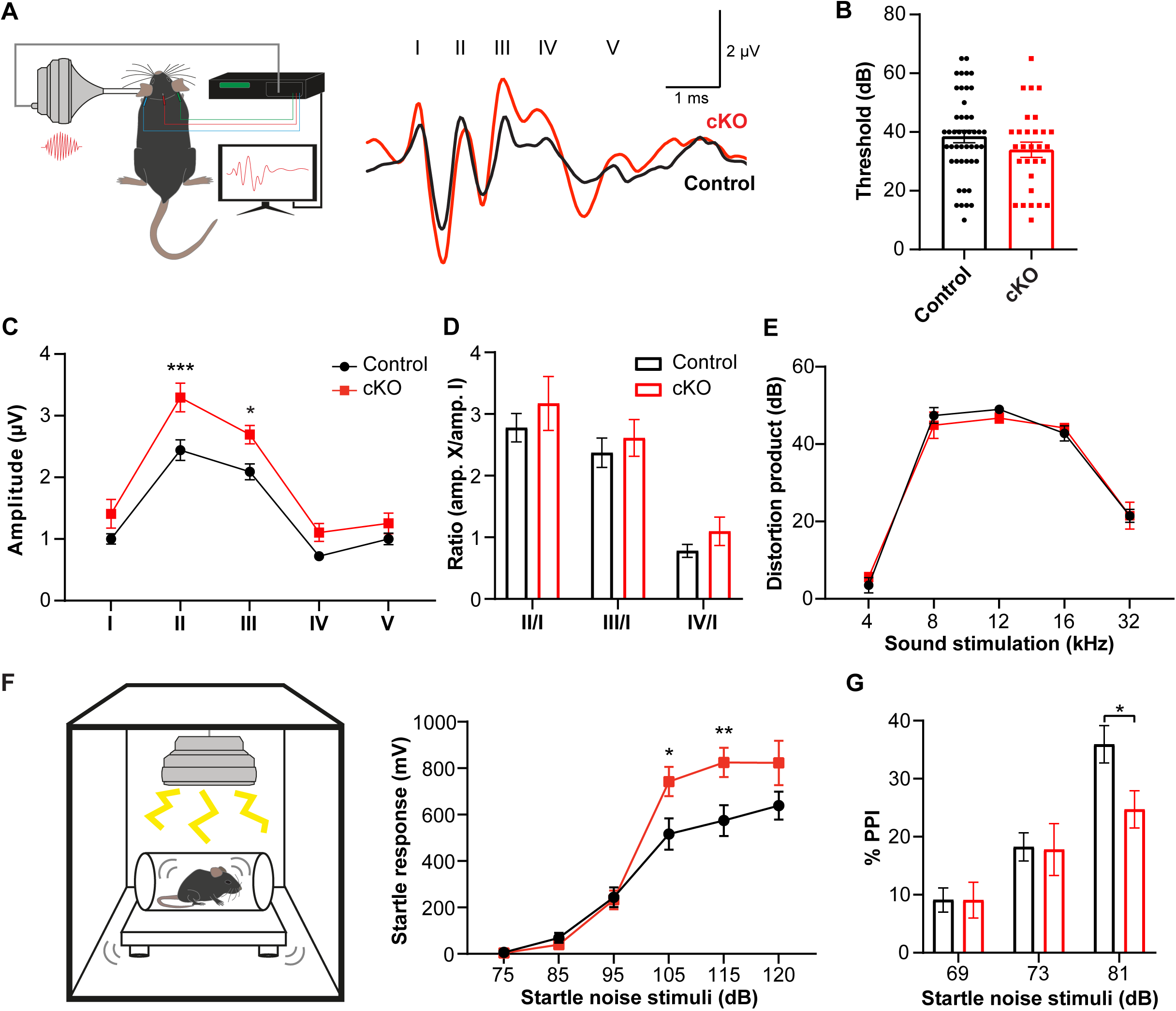
Scn2a cKO mice exhibit auditory hypersensitivity. **A.** Illustration of mouse ABR test and representative traces of ABRs in response to click stimuli (from 30 dB to 90 dB SPL) from control and *Scn2a* cKO mice at P25. **B**. Summary of ABR thresholds of control and *Scn2a* cKO mice (P21-P25) in response to click. **C.** ABR amplitudes (µV) of wave I to V responding to 80 dB SPL clicks. **D.** The ratio for the amplitude of wave I over the amplitude of each wave (II, III, and IV), evaluating central gain changes. **E.** Summary of distortion products (dB SPL) at 80 dB at various pure tone stimuli (4, 8, 12, 16, and 32 kHz). **F.** Illustration of mouse acoustic startle reflex (ASR) and summary of ASR maximum startle amplitudes (mV) to sound stimuli (from 75 dB to 120 dB SPL, background = 65 dB SPL) in control and *Scn2a* cKO mice. **G.** Pre-pulse inhibition (PPI, %) at different strength of pre-pulse stimuli (69, 73, 81 dB SPL). All values plotted are means per mouse ± s.e.m., * *p* < 0.05, ** *p* < 0.01.

To further investigate the auditory hypersensitivity in *Scn2a* cKO mice, we performed the acoustic startle reflex (ASR) test. We observed a distinct, pronounced startle response to sudden, loud auditory stimuli in *Scn2a* cKO mice, which is consistent with the hypersensitivity commonly observed in individuals with ASD(*24*). The observed startle reflexes were larger in *Scn2a* cKO mice compared to control in response to stimuli above 95 dB SPL (two-way ANOVA test, *p* = 0.0012, **Fig. 7F**). *Scn2a* cKO mice showed stronger startle at loud sound stimulation of 105 dB SPL (516.3 ± 67.6 mV, n = 15 control vs 742.5 ± 62.6 mV, n = 10 cKO, *p* = 0.0214, multiple comparison from two-way ANOVA with Bonferroni correction), 115 dB SPL (574.2 ± 66.6 mV in control vs 824.9 ± 63.1 mV in cKO, *p* = 0.0077, multiple comparison from two-way ANOVA with Bonferroni correction) and 120 dB SPL (638.5 ± 60.0 mV in control vs 822.8 ± 95.4 mV in cKO, *p* = 0.1019, multiple comparison from two-way ANOVA with Bonferroni correction). The increased amplitude of the startle reflex in *Scn2a* cKO mice indicates more sensitive at sensory perception. In addition, *Scn2a* cKO mice exhibited alterations in the pre-pulse inhibition (PPI) test. Typically, a weaker sensory stimulus (or pre-pulse) inhibits the reaction to a subsequent strong sensory event. In control, the inhibition rate drastically increased with the strength of pre-pulse stimulation. In contrast, *Scn2a* cKO mice demonstrated a diminished capacity to inhibit the startle reflex, particularly in response to the pre-pulse at 81 dB SPL sound stimulation (35.9 ± 3.22 % PPI in n = 18 control vs 24.7 ± 3.19 % PPI in n = 11 cKO mice, *p* = 0.0377, multiple comparison from two-way ANOVA with Bonferroni correction, **Fig. 7G**). The alteration in startle reflex to sound stimulation suggests an impaired sensory gating mechanism, a feature often seen in ASD (*13*). Collectively, these observations present robust behavioral phenotypes in *Scn2a* cKO mice, specifically sound hypersensitivity, and impaired sensory gating, mirroring sensory processing abnormalities in ASD. These findings underscore the pivotal role of OL-*Scn2a* mediated myelin plasticity in auditory brain circuitry, with a significant influence on sound hypersensitivity, particularly given the sensory abnormalities in *Scn2a* cKO mice.

## Discussion

Understanding the fundamental mechanisms of neuronal activity and their interplay with genetic factors provide invaluable insights for neurodevelopmental disorders, including ASD. In this study, we investigated the role of OL-*Scn2a* in the auditory nervous system in developing brain from the genetic profiling to the systemic analysis. Characterization using snRNA-seq revealed that the loss of OL-*Scn2a* impacts myelin-related gene expression in the auditory brainstem. Structural analysis of myelinated axons demonstrated thinner myelin and alterations in axons in *Scn2a* cKO mice. Electrophysiology and immunostaining showed that alterations in myelinated segments and nodal structure along the distal axon influenced the excitability of the nerve terminal, causing a reduced fidelity and reliability of spikes in *Scn2a* cKO mice. In addition, *Scn2a* cKO mice showed hypersensitivity to sound stimulation which is likely due to central gain changes. Our findings highlight the temporal progression of myelination in early development and its potential links to the onset or severity of ASD symptoms, which could provide aids to diagnosis, prognosis, and potential therapeutic interventions in the future.

### The effects of OL-*Scn2a* loss on myelin and axonal integrity

Abnormal development in white matter and myelination has been found in ASD animal models and humans with autism(*42–45*). However, how myelin changes are associated with compromised neural circuit function in ASD remains unclear. Myelination is crucial for efficient neural signal transmission. Any disruptions to the myelin sheath can influence axonal integrity, altering ion channel distribution, and consequently affect neuronal excitability(*5, 19*). Specifically, the spacing and periodicity of myelin segments determine Na_v_ channel distribution at nodes and heminodes, enabling saltatory conduction and precise neural signaling(*5, 39*). In MBP deleted rats, hypomyelination led to disorganized Na_v_ channel clustering and shorter internode, which are associated with a delayed AP onset, a longer AP half width, and AP failures at the nerve terminal(*5, 40*). In the cortical gray matter neuronal circuitry, cuprizone-induced demyelination caused hyperexcitability in pyramidal neurons by altering the axon initial segment (AIS) position and reducing the efficacy of AP generation(*19*). The current study expands the dimensions of what have been previously reported by examining the effects of OL-*Scn2a* loss on myelin and axonal integrity specifically at distal axons near nerve terminals. We demonstrate that OL-*Scn2a* loss leads to alterations in ion channel redistribution and aberrant excitability at nerve terminals such as conduction failure and spontaneous spiking. This is particularly pertinent as *Scn2a*-expressing immature OLs are abundant in synapse-rich regions like the MNTB. Interestingly, we observed that OL-*Scn2a* loss had more pronounced effects at nerve terminals than on nodes, which differs from prior models like MBP-deleted or cuprizone-induced demyelination. In these models, mature OLs were either unable to myelinate properly or underwent cell death(*5, 19*). In contrast, *Scn2a* cKO mice may have impaired interactions between immature OLs and distal axons, significantly affecting nerve terminal function during postnatal development. Therefore, our study underscores the region-specific relationship between myelin integrity and ion channel distribution in the developing brain. We emphasize that any disturbances in myelin structure can trigger cascading effects on neuronal excitability and synaptic function in the CNS, especially at nerve terminals in the auditory nervous system.

### Role of *Scn2a* in OL development and myelination

While OL dysfunctions are highly associated with the pathophysiological process of ASD, few studies have focused on ion channels of OLs. Our findings highlight the roles of *Scn2a*, a gene highly linked to ASD, in OL development and OL interaction with axons. Given that mature OLs play an indispensable role in forming the myelin sheath understanding how *Scn2a* is involved in their maturation is important. A variety of ion channels, including Na_v_, K_v_, and voltage-gated calcium channels (Ca_v_), as well as inward-rectifier potassium channels (K_ir_), are expressed in OLs(*46, 47*). While the functions of Ca_v_ and K_ir_ channels in OLs are relatively well-defined, the roles of Na_v_ remain unclear. Previous studies spotlighted the expression of Na_v_1.2 in immature OLs with a spiking phenotype and highlighted the potential role in OL differentiation(*26, 27*). Supporting this, our snRNA-seq data demonstrated a decline in the mature OL population coupled with an increase in OPCs and differentiating OLs in *Scn2a* cKO mice. In addition, OL*-Scn2a* deletion was found to down-regulate myelin-related genes in mature OLs. How are Na_v_1.2 channels, encoded by *Scn2a*, involved in OL maturation and myelination? One possible explanation is that the activation of Na_v_1.2 may be pivotal for triggering Ca_v_ channel activation, leading to a Ca^2+^ flux within OLs, which is involved in OL proliferation, migration, and differentiation(*47*). Specifically, Ca^2+^ signaling facilitated by R-type Ca_v_ in myelin sheaths at paranodal regions, might influence the growth of myelin sheaths(*46, 48*). To activate high-voltage activated calcium channels such as L- and R-Type efficiently, the activation of Na_v_1.2 channels should be required for depolarizing OL membrane to around -30 mV (*49*). Consequently, the synergic interplay between Na_v_1.2 and Ca_v_ channels could amplify calcium signaling in OLs, initiating the differentiation and maturation processes(*47*).

### Auditory hypersensitivity in *Scn2a* cKO mice

One prevalent characteristic of ASD is sensory processing disorders, notably auditory hypersensitivity. Interestingly, in various autistic mouse models, the outcomes from auditory functional tests, particularly ABRs, have been inconsistent. For example, C*ntnap2* KO mice exhibited normal hearing thresholds, but with prolonged latencies of ABRs. *Fmrp1* KO mice showed hypersensitivity in hearing perception attributed to aberrant activity in the auditory cortex (*50, 51*). Similarly, *Shank3* KO mice displayed auditory hypersensitivity, as evidenced by amplified ABR responses and a heightened startle reflex in the ASR(*52*). In the current study, *Scn2a* cKO mice showed an augmented amplitude of ABRs and a stronger startle reflex, despite no peripheral changes. The results lead to the question: How are alterations in myelin and neuronal properties associated with auditory hypersensitivity in *Scn2a* cKO mice? The auditory system, with its intricate neural pathways and high demand for precise timing, is particularly vulnerable to disruptions in signal conduction. Defects in myelination within auditory pathways can lead to synchronization issues among neurons responsible for sound processing. Increased asynchronized signals can compromise the precise timing required for sound localization and the discernment of complex auditory patterns. In *Scn2a* cKO mice, a loss of temporal fidelity at presynaptic terminals and inconsistent spike conduction along myelinated axons was observed. Notably, ∼20% of presynaptic recordings exhibited aberrant and asynchronous spikes at the nerve terminal in *Scn2a* cKO. The atypical firing of auditory neurons can contribute to auditory hypersensitivity. Neuronal firing rates were abnormally increased following demyelination, resulting in hyperexcitability of the neural circuitry(*19, 53*). This hyperexcitability may be caused by alterations in sodium channel distribution and a rise in extracellular potassium along the myelinated axon(*54*). Another possibility is that hypersensitivity may also emerge from changes in the intricate balance between excitatory and inhibitory regulation within the auditory brainstem. A decline in inhibitory neurons or disruptions in inhibitory inputs can induce hypersensitivity(*55, 56*). In *Shank3* KO mice, hypersensitivity might be caused by diminished inhibitory regulation in the auditory circuit(*52*). In addition, our snRNA-seq data suggested a downward trend in genes associated with neuronal migration in interneuron populations from *Scn2a* cKO mice (data not shown). In conclusion, the integrity of the myelin sheath plays a pivotal role in regulating neuronal excitability and ensuring proper auditory function. Defects in myelination can create a spectrum of auditory dysfunctions, including hypersensitivity. Our results demonstrated how OL*-Scn2a* is involved in the relationship between myelin defects, neuronal excitability, and auditory pathology in ASD, potentially paving the way for targeted therapeutic interventions.

## Materials and Methods

### Animals

All procedures were approved in advance by the Institutional Animal Care and Use Committee of University of Michigan. To create OL-specific *Scn2a* knockout mice, we crossed a *Scn2a*^fl/fl^ mice with *Pdgfra*^CreERT^ mice, generating double transgenic mice (*Pdgfra*^CreERT^; *Scn2a*^fl/fl^) as described in a previous study (*27*). Littermates without a Cre-recombinase but with *Scn2a*^fl/fl^ were used as control. 70 mg/kg of tamoxifen was administered via i.p. injection at postnatal days (P) 4, 6, and 8. Both male and female mice aged P10-P17 were used for EM, immunohistochemistry, and ex vivo electrophysiology. Mice of both sexes aged P21-P27 were used for snRNA-seq, ABR, ASR, and DPOAE.

### Slice preparation

After rapid decapitation of the mice, which were deeply anesthetized by isoflurane inhalation, the brainstem was quickly removed from the skull and immersed in ice-cold low-calcium artificial CSF (aCSF) containing the following (in mM): 125 NaCl, 2.5 KCl, 3 MgCl_2_, 0.1 CaCl_2_, 25 glucose, 25 NaHCO_3_, 1.25 NaH_2_PO_4_, 0.4 ascorbic acid, 3 myoinositol, and 2 Na-pyruvate, pH 7.3–7.4 when bubbled with carbogen (95% O_2_/5% CO_2_), and 310–320 mOsm/L. The brainstem was sectioned (200 μm thick for electrophysiological recordings and immunostaining) and the slices were transferred to an incubation chamber containing normal aCSF bubbled with carbogen, where they were maintained for 30 min at 34-35°C and thereafter at room temperature (24°C). Normal aCSF was the same as low calcium aCSF, but with 1 mM MgCl_2_ and 2 mM CaCl_2_.

### Single nuclei RNA sequencing (snRNA-seq)

A snRNA-seq was conducted using brainstem tissues from control (*Scn2a*^fl/fl^) and *Scn2a* cKO mice (*Pdgfra*^CreERT^; *Scn2a*^fl/fl^). The brainstem was dissected and mechanically homogenized using dounce homogenizer in homogenizing buffer (250 mM sucrose, 25 mM KCl, 5 mM MgCl_2_, 10 mM Tris, 1 uM DTT and 0.1% Triton-X100) supplemented with enzymatic RNase Inhibitor (400 U/ml). 700 µl of homogenization solution was added. Homogenization involved 5 strokes of a loose pestle and 10-15 strokes of a tight one. The resultant solution was completed to 1 ml and passed through a 40μm strainer. Post centrifugation, nuclei were rinsed with PBS containing RNAse inhibitor. A subsequent filtration through a 20 µm strainer was done before resuspending in 1 ml of PBS fortified with RNAse inhibitor. For fixation, 3 ml of 1.33% PFA was added to the nuclei for 10 minutes, followed by permeabilization using 160 μL of 5% Triton X-100 for 3 minutes. After washing out the PFA, nuclei were suspended in PBS with RNAse inhibitor and quantified manually via a hemocytometer. The fixed nuclei were barcoded and prepared for library using Evercode WT Mini kit (Parse Biosciences) following manufacturer’s guidelines. The final library underwent sequencing on Novaseq 500 at Novegen (Sacramento, CA).

The raw reads were mapped and quantified using *split-pipe* v1.0.3 (Parse Biosciences). The count data was analyzed using *Seurat* (*57*) in R. To minimize artificial error, the cells with more than 1% mitochondria DNA and 5000 transcripts were considered dead cells and doublet cell, and those were excluded from further analysis. All nuclei were clustered using shared nearest neighbor (SNN) and plotted using *Seurat* in R. Each cluster was manually identified by examining expression of cell markers (*Pdgfra/Cspg4* for OPC, *Mog/Mbp/Plp1* for mature OL, *Aldh1l1* for astrocyte, *Itgam/Csf1r* for microglia, *Pecam1* for vascular cells, *Rbfox3* for neuronal population). To examine oligodendrocyte differentiation, the oligodendrocyte lineage clusters were isolated from the data set and trajectory analysis was conducted using *Monocle3* in R (*58*). The population of each differentiation stage of oligodendrocytes was calculated from [number of nuclei from each cluster/total number of nuclei from all of OL lineage]. To examine gene expression pattern in each cluster, the differentially expressed genes between control and cKO were obtained with *DESeq2* (*59*) method with Bonferroni multiple comparison correction in *Seurat*. For further analysis, we isolated matured OL cluster, where showed significant gene expression difference between groups, and examined expression level of myelin related genes. To list myelin related genes, Amigo2 (*60*) was used. 391 genes were searched on Amigo2 using the keyword, “myelin”. The expression level of the genes was compared between groups using *DESeq2* methods without multiple correction. The DEGs from the analysis were used gene network analysis using String (*61*). The network was manually categorized based on their function. All of data visualization was done using *ggplot2* (*62*)and String.

### Electrophysiology

Slices were perfused with normal aCSF at 2 ml/min and visualized using an infrared differential interference contrast microscope (AxoExaminer, Zeiss, Oberkochen Germany) with a 63× water-immersion objective and a CMOS camera (ORCA-Flash2.8, Hamamatsu, Japan). Whole-cell patch-clamp recordings were performed in normal aCSF at room temperature (24°C) using an EPC-10 amplifier controlled by PATCHMASTER software (HEKA, Elektronik, Lambrecht/Pfalz, Germany). For presynaptic recordings, the pipette solution contained (in mM): 125 K-gluconate, 20 KCl, 5 Na_2_-phosphocreatine, 10 HEPES, 4 Mg-ATP, 0.2 EGTA, and 0.3 GTP, pH adjusted to 7.3 with KOH. Recordings were not corrected for the predicted liquid junction potential of 11 mV. Patch electrodes had resistances of 4–5 MΩ. Current-clamp recordings were continued only if the initial uncompensated series resistance was <20 MΩ (*4, 5*). Lucifer Yellow (1 mM, Invitrogen) was added to the pipette solution to visualize the calyx of Held terminal. Presynaptic action potentials (APs) from the calyx of Held terminal were evoked by stimulation with a bipolar platinum-iridium electrode (Frederick Haer, Bowdoinham, ME) placed near the midline spanning the afferent fiber tract of the MNTB. An Iso-Flex stimulator driven by a Master 10 pulse generator (A.M.P.I., Jerusalem, Israel) delivered 100-µs pulses at 1.2 times threshold (<15 V constant voltage). Signals were filtered at 2.9 kHz and acquired at a sampling rate of 10–50 µs. AP waveform parameters were analyzed from the first AP induced by minimum current injection (rheobase current) and the subsequent AP phase plot, where membrane potential slop (dV/dt) is plotted against the membrane potential(*63*). In both and *Scn2a* cKO mice, all cells displayed a single inflection in the rising phase of the AP, indicating APs were generated at the heminode adjacent to the presynaptic terminal. Presynaptic AP trains were obtained by averaging three sweeps (five for a single AP) in each experiment. Data were analyzed offline and presented using Igor Pro (Wavemetrics, Lake Oswego, OR).

### Auditory brainstem responses (ABRs)

To record ABR, the mice (P21-P27) were anesthetized with 3.5% isoflurane and maintained with 2.3% isoflurane during recording (1 l/min O_2_ flow rate). ABR recordings were performed in a sound attenuation chamber (Med Associates, Albans, VT). Subdermal needle electrodes (Rochester Electro-Medical, Lutz, FL) were placed on the top of the head, ipsilateral mastoid, and contralateral mastoid as the active, reference, and ground electrode, respectively. The signal differences in the ABRs between the vertex and the mastoid electrodes were amplified and filtered (100–5000 Hz). Acoustic stimuli were generated by an Auditory Evoked Potentials Workstation (Tucker-Davis Technologies [TDT], Alachua, FL). Closed field click stimuli were presented to the left ear. The signals consisted of a series of amplitude-modulated square waves (0.1 ms duration, 16/s) through TDT Multi-Field Magnetic Speakers. The sound stimuli were delivered through a 10-cm plastic tube (Tygon; 3.2-mm outer diameter) at a repeat rate of 16/s. Sound intensities ranged from 90 to 20 dB, with 5-dB decrements, and responses to 512 sweeps were averaged.

### Distortion Product Otoacoustic Emissions (DPOAEs)

Mice were anesthetized with 3.5% isoflurane and maintained with 2.3% isoflurane during recording. DPOAE recordings were performed in a sound attenuation chamber (Med Associates, Albans, VT). Acoustic stimuli were generated by an Auditory Evoked Potentials Workstation (TDT, Alachua, FLO). The ER-10B+ recording microphone (Etymotic Research, Elk Grove Village, IL) with ear-tip was inserted into the ear canal. The sound stimuli were delivered through two TDT Multi-Field Magnetic Speakers connected to the recording microphone by 10-cm coupling tubes (Tygon; 3.2 mm outer diameter). Pure tones were presented at 20% frequency separation between f1 and f2 at 8, 12, 16, and 32 kHz. Sound intensities ranged from 80 to 20 dB, with 10-dB decrements, and responses to 512 sweeps were averaged. Distortion products were calculated as 2f1-f2 minus the noise floor that were detected by the recording microphone and amplified by RZ6 processor (TDT).

### Acoustic startle responses (ASRs)

Mice of either sex between ages P25-P27 were put in a Plexiglas holding cylinder located in sound-attenuated chamber using the SR-LAB startle response system (San Diego Instruments, San Diego, CA). Weights of mice were recorded to account for sex or size differences. Sound levels from the chamber speakers were calibrated with a digital sound level meter (Part Number MS-M80A, Mengshen). Each trial consisted of an initial acclimation period of 5 minutes of background level noise (at 65 dB) followed by 5 rounds of randomized noise stimuli playing for 40 milliseconds with 15 seconds of background between each noise stimuli at 65 dB (background), 95 dB, 105 dB, 115 dB, and 120 dB. Stimuli was produced by a digital signal processing-controlled system amplified and emitted by a loudspeaker. Startle responses were measured inside the sound-attenuated chamber by a movement-sensitive piezo-accelerometer platform. The maximum was used as the startle amplitude at each tone with millivolts (mV) as the unit of measurement and averaged per mouse. For each strength of stimulus, startle amplitudes were averaged across 10 trials. Startle responses to the three initial stimuli were excluded from statistical analyses.

### Pre-pulse inhibition (PPI) of acoustic startle response

PPI of ASR was examined 2 days after the ASR assessment. The apparatus and basic experimental conditions were identical to that described above. Test sessions started with the acclimatization period which included three startling stimuli (120 dB) to accustom the mice to the experimental procedure. The initial stimuli were followed by 100 trials (10 × 10 trials) presented in a random order. The PPI session involved: 10 trials with a sham stimulus (65 dB, 40 ms), three types (3 × 10) of pre-pulse trials (PP) which included only 20 ms PP stimuli (69, 73, and 81 dB), 10 pulse trials (P) which included only a pulse (startling) stimulus (120 dB, 40 ms), three types (3 × 10) of pre-pulse-and-pulse trials (PP-P) which included a 20 ms PP (69, 73, and 81 dB) followed 100 ms later by a 120-dB P stimulus. Startle responses were measured for 100 ms after the onset of the last stimulus within each trial. For each type of trial, startle amplitudes were averaged across 10 trials. The magnitude of PPI was calculated as a percent inhibition of the startle amplitude in the P trial (treated as 100%) according to the formula: [startle amplitude in P trials – startle amplitude in PP-P trials)/startle amplitude in P trials] × 100%. Startle responses to the three initial stimuli were excluded from statistical analyses.

### Immunohistochemistry

Mouse brainstem slices (200 µm) stained with tetramethylrhodamine dextran (Invitrogen) were incubated in normal aCSF bubbled with carbogen at 37°C for 30 min. All slices were fixed with 4% (w/v) paraformaldehyde in PBS for 10 min. Free-floating sections were blocked in 3% goat serum and 0.3% Triton X-100 in PBS for 30 min and incubated with the primary antibody overnight at room temperature. The following primary antibodies were used: anti-PanNa (mouse IgG1; 1:400; Sigma, Cat. #S8809), anti-Caspr (guinea pig IgG; 1:200; gifted from Dr. Manzoor Bhat, UTHSCSA), anti-PLP1 (mouse IgG2a, 1:500; invitrogen, Cat. #MA1-80034). Antibody labeling was visualized by incubation of appropriate Alexa fluor–conjugated secondary antibodies (1:500; Invitrogen) for 2 h at room temperature. Stained slices were viewed with laser lines at 488 nm, 568 nm, and 647 nm using a 40x/1.40 or 63ξ/1.40 oil-immersion objective on a confocal laser-scanning microscope (LSM-710; Zeiss). Stack images were acquired at a digital size of 1024 × 1024 pixels with optical section separation (*z*-interval) of 0.5 µm and were later crop to the relevant part of the field without changing the resolution. The confocal image stacks were analyzed using ImageJ software.

### Transmission electron microscopy

The brain was removed and immersed in ice-cold low-calcium aCSF (mentioned previously in slice preparation) and the MNTB from the brainstem was dissected at 200 µm-thick section using a microtome (VT1200s, Leica), then fixed with fixative consisting of 4% formaldehyde and 1% glutaraldehyde and stored at 4°C. Further processing was performed by the UTHSCSA Electron Microscopy Lab as previously described (*64, 65*). Axon bundle images were analyzed at a final magnification of 8000X, with the longest length of the axon measured as the inner diameter and the inner radius divided by the outer radius as the *g*-ratio. Three images of axon bundles per mouse were analyzed and averaged.

### Statistical analysis

Immunostaining data were based on analyses from at least six cells in six slices from three to eleven animals. Experimental data were analyzed and presented using Igor Pro and Prism (GraphPad Software, San Diego, CA). For statistical significance, we tested the normality of the data distribution with the Kolmogorov-Smirnov test with the Dallal-Wilkinson-Lillie *p*-values using Prism 5. If a dataset passed the normality test, we used the parametric test (unpaired *t*-test for two group comparison, ANOVA for multiple comparison); for all other datasets we used the non-parametric test (Mann-Whiney U test for two group comparison). For linear regression analysis slope and intercept values were compared. Data collected as raw values are shown as mean ± S.E.M. Details of statistical methods are reported in the text. For all analyses, *p*-values < 0.05 were considered significant.

## Acknowledgments

We are grateful to Dr. Manzoor Bhat (UTHSCSA) for providing reagents, insightful discussion, and technical assistance. We also thank Drs. Kevin Bender (UCSF) and Shin Kang (Temple) for providing transgenic mice. We thank Drs. Jason Pugh and Michael Roberts for providing comments on the manuscript.

## Author contributions

Conceptualization: JHK

Methodology: HGB, WCW, KN, EG, JHK

Investigation: HGB, WCW, KN, EG, JHK

Visualization: HGB, WCW, KN, JHK

Supervision: JHK

Writing—original draft: HGB, WCW, JHK

Writing—review & editing: HGB, WCW, JHK

## Data and materials availability

Raw genetic data can be accessed on GEO NCBI with the accession number GSE252185, and analysis script can be accessible as supplementary data. All other data are available in the main text or the supplementary materials.

## Supplementary Materials

**Fig. S1. Expression patterns of cell specific markers in snRNA-seq. A-F.** To identify cell types of the clusters, the expression patterns of cell markers were examined. The expression patterns of cell specific markers were plotted on UMAP plot: *Cspg4* for OPC (**A**), *Plp1* for mature OL (**B**), *Aldh1l1* for astrocyte (**C**), *Itgam* (**D**) and *Csf1r* for microglia (**E**), *Pecam1* for epithelia cell (**F**), *Rbfox3* for neuron (**G**), *Gad1* for inhibitory neuron (**H**), and *Slc17a7* for glutamatergic neuron (**I**). Scaled expression level was color coded on the side of each plot.

**Fig. S2. Axon caliber and myelin thickness deficits from *Scn2a* cKO mice. A.** Representative TEM image of axons in the MNTB region from a different OL-specific cKO mouse line (*Sox10*^CreER^; *Scn2a*^fl/fl^, at P25) with an inducible *Sox10* Cre (*Sox10*^CreER^) targeting all OL lineage cells. **B.** Scatter plot of *g*-ratio values plotted as a function of corresponding axon inner diameter of individual axons in the MNTB region. Datapoints from individual axons and best fit lines are shown in black for control and purple for *Scn2a* cKO mice (*Sox10*^CreER^; *Scn2a*^fl/fl^). **C.** Summary of inner diameter of individual axons (left), their respective cumulative frequency (middle), and the histogram distribution of inner diameter (right). **D.** Summary of *g*-ratio of individual axons (left), their respective cumulative frequency (middle), and the histogram of distribution of *g*-ratio (right). Note that *Scn2a* cKO mice exhibit a higher *g*-ratio and larger inner diameter of axons in the auditory brainstem, indicating alterations in myelination and axonal integrity. Values are shown as mean ± s.e.m., *** *p* < 0.001, **** *p* < 0.0001.

**Fig. S3. Conduction failures at the calyx terminals from *Scn2a* cKO mice. A.** Representative traces of presynaptic APs from control (black) and *Scn2a* cKO mice (*Sox10*^CreER^; *Scn2a*^fl/fl^, purple) stimulated at 200 Hz. Grey arrowheads indicate absent APs. **B.** Left, violin plots showing AP failure rate from control and *Scn2a* cKO mice stimulated at varied frequencies. Solid and dashed lines represent median and first and third quartile, respectively. Right, average AP failure rate at varied stimulating frequencies. **C.** Average threshold intensity (TI) to initiate a single AP from control and *Scn2a* cKO mice. Individual datapoints are displayed as circles and triangles. **D.** Representative traces of 200 Hz AP train from *Scn2a* cKO mice stimulated with TI, 20% increment from TI, and 40% increment from TI. **E.** Left, 200 Hz AP failure rate of individual cells from incremental stimulating intensities. Right, average 200 Hz AP failure rate from incremental stimulating intensities (mean ± s.e.m.). * *p* < 0.05, *** *p* < 0.001.

**Fig. S4. Aberrant spikes and spontaneous firing are present in some calyx terminals from *Scn2a* cKO mice. A.** Left, representative trace of aberrant spikes at the calyx terminal. In *Scn2a* cKO mice (*Pdgfra*^CreERT^; *Scn2a*^fl/fl^), APs were evoked by afferent fiber stimulus with high temporal fidelity but followed by aberrant spikes. Right, ∼25% of calyces (6/26 cells in *Scn2a* cKO) exhibited this abnormal spiking. However, all recordings from control did not show any aberrant spikes or spontaneous firing (0/20 cells in control). Arrow indicates the stimulus artifact caused by afferent fiber stimulation. **B.** Left, representative trace for spontaneous firing. The calyces (∼25%) displayed spontaneous firing around -65 mV with a bursting rate of 200 Hz. Right, expanded view of the burst spiking in box.

**Table S1 DE analysis from OPC**

**Table S2 DE analysis from Differentiating OL 1**

**Table S3 DE analysis from Differentiating OL 2**

**Table S4 DE analysis from premature OL**

**Table S5 DE analysis from mature OL**

