## Supplementary figures and images for "*Scn2a* deletion disrupts oligodendroglia function: Implication for myelination, neural circuitry, and auditory hypersensitivity in ASD"

### Fig. S1

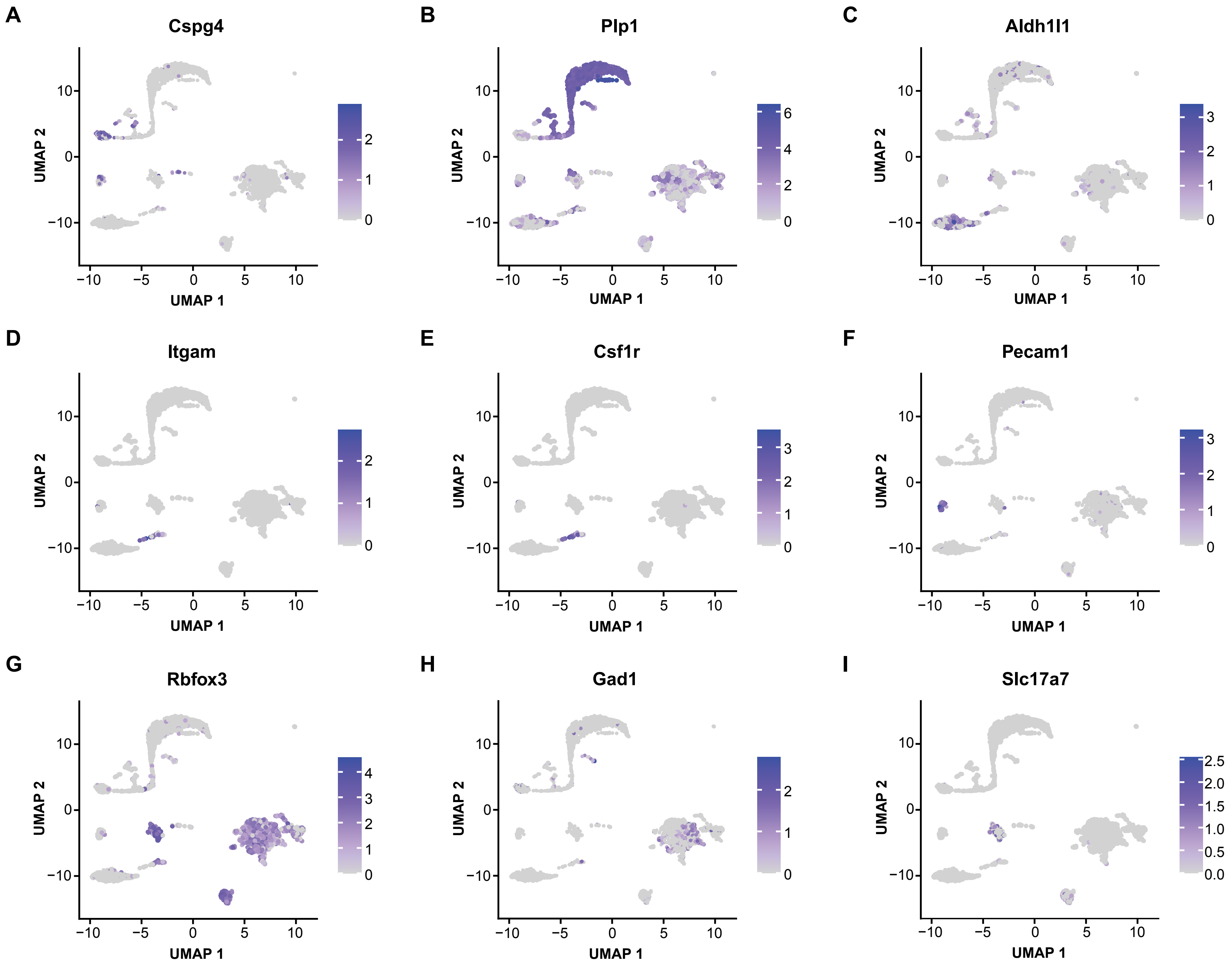

### Fig. S2

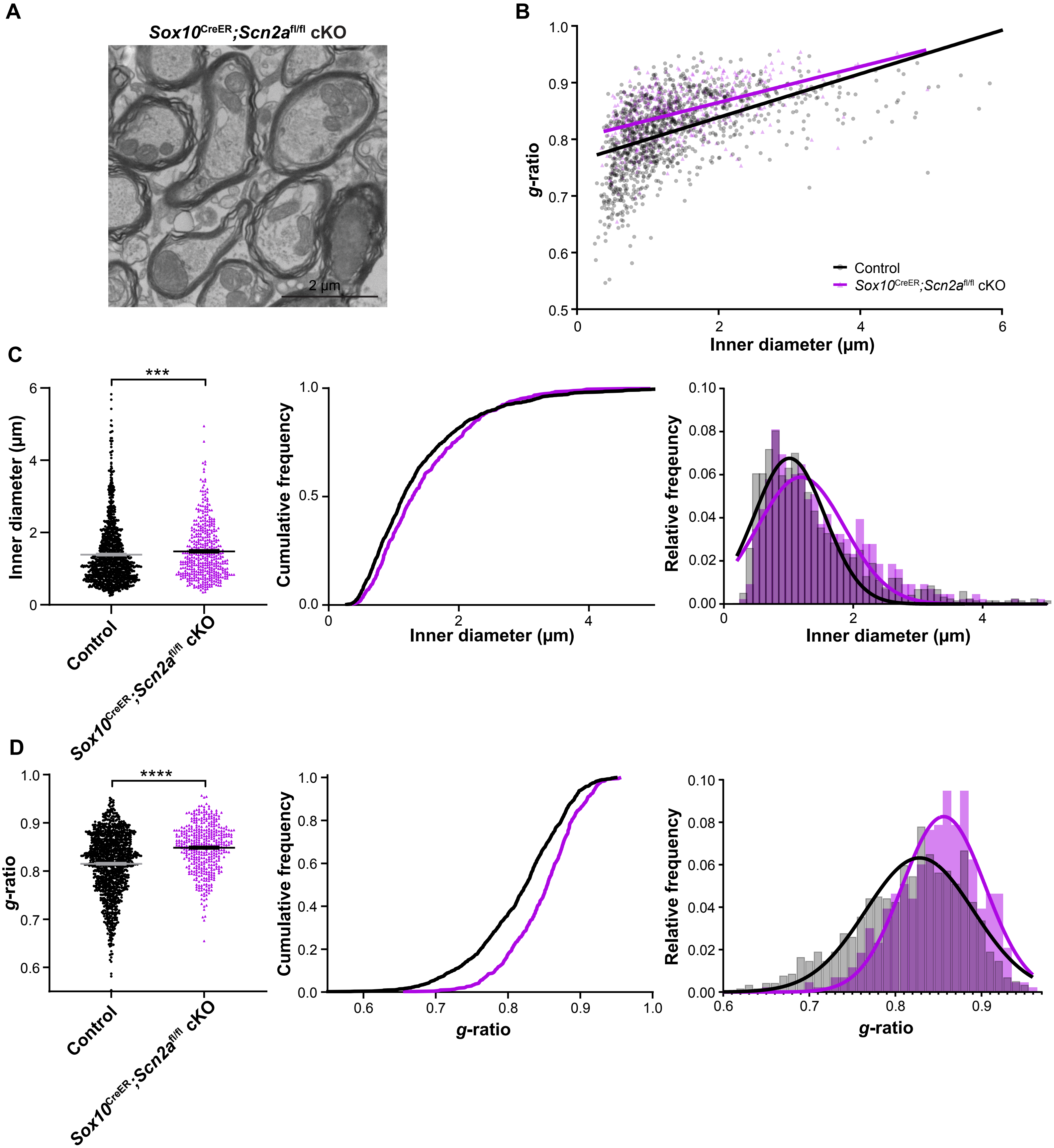

### Fig. S3

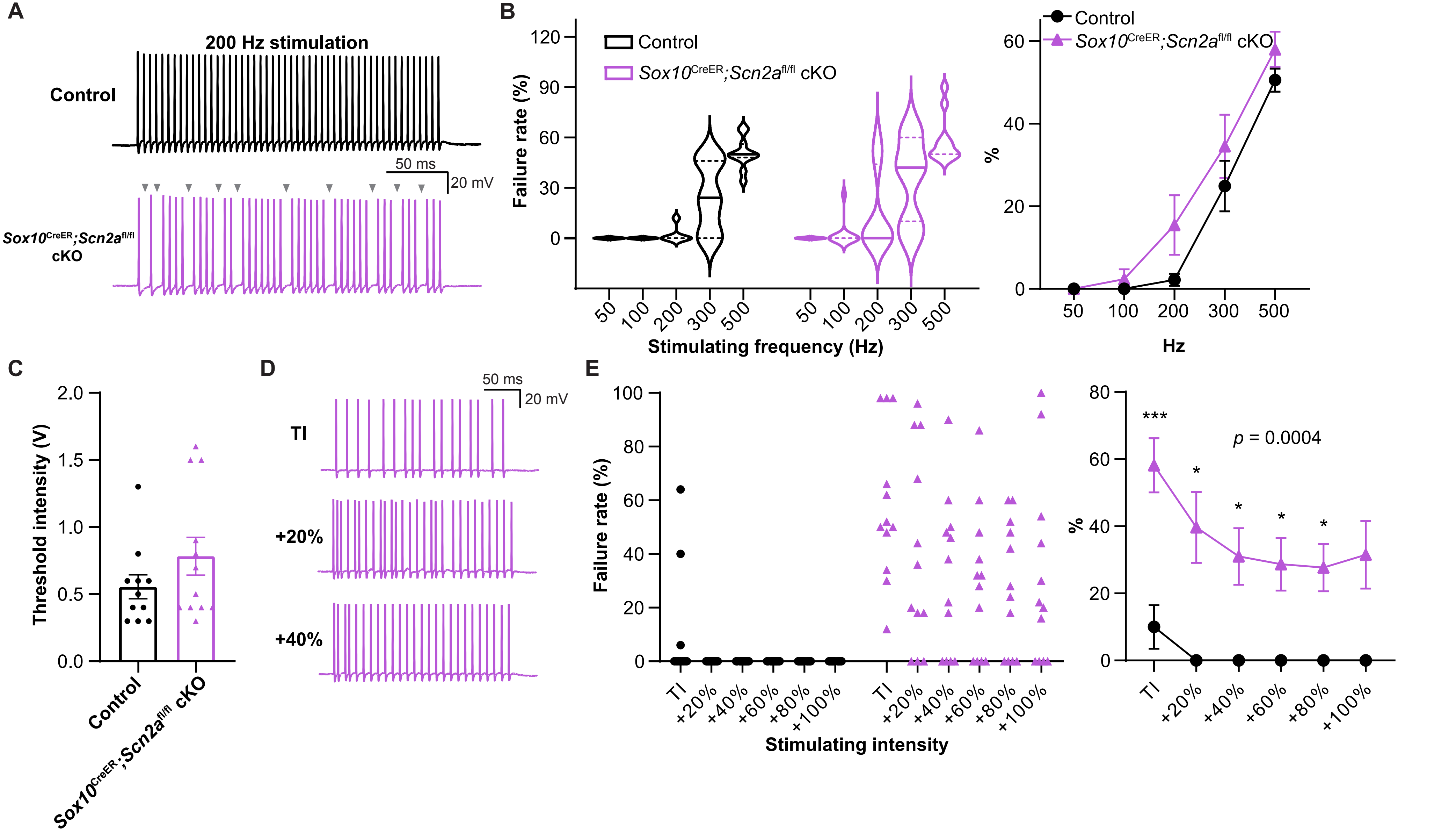

### Fig. S4

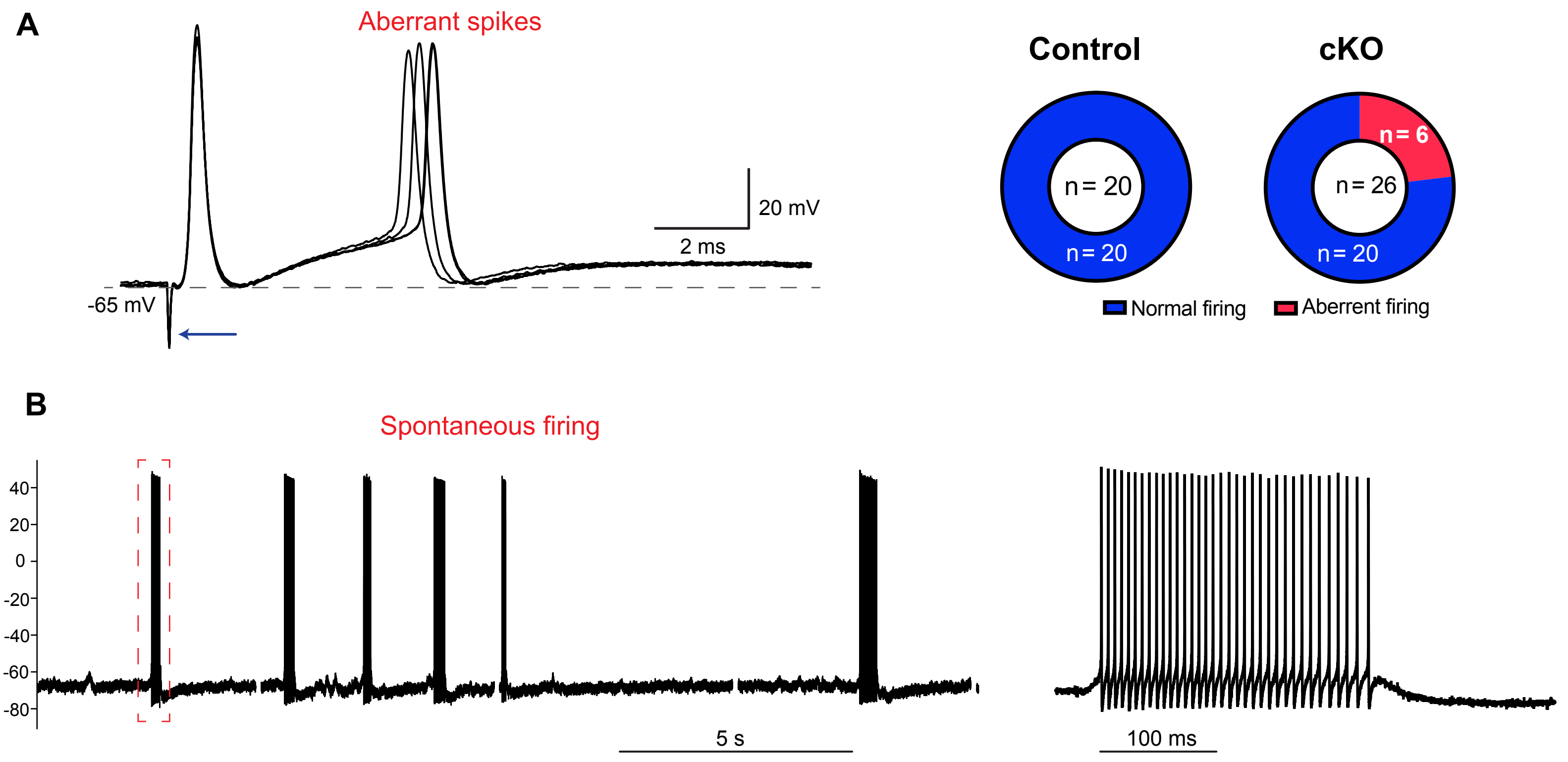
